# Dual Function of Mitochondrial Complex III in *Plasmodium falciparum*

**DOI:** 10.1101/2025.10.02.680181

**Authors:** River S. Rell, Anurag Shukla, Joanne M. Morrisey, Ijeoma C. Okoye, Michael W. Mather, Akhil B. Vaidya

**Affiliations:** Center for Molecular Parasitology, Department of Microbiology and Immunology, Drexel University College of Medicine, Philadelphia, PA 19129

## Abstract

Complex III of the malaria parasite mitochondrial electron transport chain (mtETC) has been validated as an attractive target for currently used antimalarials. We previously showed that the main function of mtETC in blood stage *Plasmodium falciparum* is to regenerate ubiquinone, which serves as an obligatory co-substrate of dihydroorotate dehydrogenase (DHOD), an essential mitochondrial enzyme for pyrimidine biosynthesis. *P. falciparum* can be rendered resistant to all mtETC inhibitors by provision of a bypass mediated by cytosolic yeast DHOD, a fumarate-reducing enzyme. Malaria parasite mitochondrial DNA (mtDNA) encodes only 3 proteins, each a component of mtETC. However, attempts to eliminate mtDNA in transgenic parasites expressing yDHOD have been unsuccessful, suggesting the possibility that essential function(s) other than the canonical redox reactions of the mtETC also require mtDNA maintenance. Here we have tested the hypothesis that Complex III serves the dual functions of processing imported mitochondrial proteins, as well as ubiquinone regeneration. We have generated transgenic lines that conditionally express mitochondrial processing peptidase a (MPPα), which is also a component of Complex III. Using these parasites, we have determined that MPPα is essential even when the need for mitochondrial electron transport is bypassed. MPPα knockdown also resulted in hypersensitivity of the parasites to proguanil, a drug that synergizes with mtETC inhibitors such as atovaquone. Pulldown with MPP*α* followed by proteomics revealed the association of multiple mitochondrially targeted proteins, in addition to all components of Complex III. These results are consistent with the suggestion that Complex III in *P. falciparum* serves both mtETC and protein processing functions in mitochondrial physiology.

## Introduction

Mitochondrial Complex III in Apicomplexan parasites is a clinically validated drug target, and inhibitors of it are in therapeutic use. Divergence of the parasite Complex III from the host underlies the selectivity of these potent drugs (1, 2). Working through a modified Q cycle (3, 4), Complex III catalyzes the transfer of electrons from ubiquinol to cytochrome *c* while transporting protons into the mitochondrial intermembrane space. Inhibition of this step in mtETC results in blockage of all mitochondrial dehydrogenases that require ubiquinone as the electron acceptor. In previous studies, we have shown that in the blood stage *P. falciparum* the only critical dehydrogenase for parasite survival is dihydroorotate dehydrogenase (DHOD), an essential enzyme that is required for pyrimidine biosynthesis by the parasites (5). When the parasites are genetically modified to express cytosolic yeast DHOD (yDHOD), they are rendered independent of mtETC, since yDHOD uses fumarate as the electron acceptor, rather than ubiquinone (5). This metabolic bypass rendered the parasites resistant to all mtETC inhibitors such as atovaquone. Mitochondrial DNA (mtDNA) in malaria parasites is highly reduced in size and encodes only three proteins, each a component of mtETC (cytochrome *b* and subunits I and III of cytochrome c oxidase) (1, 6, 7). Therefore, we reasoned that it should be possible to eliminate mtDNA from mtETC-independent parasites expressing yDHOD. However, our multiple attempts to achieve this by disrupting genes encoding mitochondrial functions required for mtDNA maintenance and mitochondrial translation have failed in transgenic parasites expressing yDHOD (8-10). Thus, it is likely that the mtETC complexes serve one or more functions in addition to respiration in blood stage malaria parasites. At present the nature of these functions remains unknown.

Except for the 3 proteins encoded by the parasite mtDNA, all mitochondrial proteins are encoded in the nuclear genome, translated in the cytoplasm and imported into the mitochondrion. Many of these proteins contain targeting peptides at their N-termini. These peptides are cleaved off by mitochondrial processing peptidase (MPP) (11, 12). MPPs in mammalian and yeast mitochondria consist of heterodimers of α and β subunits that are localized within the mitochondrial matrix (13, 14). The catalytic sites are present within the β subunit, which is a metalloprotease with a Zn^2+^ ion at the active site (15). The α subunit does not have catalytic property but is essential for substrate recognition and proper presentation of the peptide within the catalytic site in multiple steps (16, 17). In addition to the matrix-localized MPP in fungi and mammals, Complex III in eukaryotes also contain Core1 and Core2 subunits that have high sequence homology to the MPP subunits α and B, respectively (17-19). These subunits in yeast have lost their catalytic activity, whereas in mammals they serve to process just one protein, the Rieske 2Fe2S subunit of Complex III (20, 21). Hence, the majority of mitochondrial proteins in these organisms have to be processed by the matrix located MPP. In stark contrast, plant mitochondria do not contain matrix located MPPs, which led Braun to suggest that Complex III associated Core proteins fulfilled MPP function, as later confirmed, and thus Complex III has dual functions in plants: ubiquinol oxidation to serve mtETC and processing of mitochondrially targeted proteins (22, 23). We hypothesize that in malaria parasites correct assembly of Complex III is essential not just for its function in mtETC but also for processing most imported mitochondrial proteins. Thus, even when mtETC has been rendered redundant, Complex III would still need to be assembled to process mitochondrial proteins. In this study we have used conditional expression of MPPα to investigate the potential dual role of Complex III in blood stage *P. falciparum*.

## Materials and Methods

### *P. falciparum* culture and transfection

We used *P. falciparum* D10_VP2KO, in which the pfVP2 locus encoding a type 2 H^+^ pumping pyrophosphatase was disrupted with the yDHOD selectable maker, for transfections (24). These parasites are resistant to all mtETC inhibitors, as well as inhibitors of parasite DHOD. *D10_VP2KO and Dd2attb* asexual blood stage parasites were cultured in human (O+) RBC, at 2.5% hematocrit in RPMI1640 medium supplemented with 0.5% Albumax, 2 g/liter NaHCO_3_, 15 mM HEPES pH 7.4, 1 mM hypoxanthine and 50 mg/liter gentamicin in 90% N_2_, 5% O_2_ and 5% CO_2_. Testing for Mycoplasma was done on a regular basis to ensure the cultures were free of any such infection.

For transfection, parasite cultures with 5-7% ring stage parasites were washed three times with Cytomix. The parasites were resuspended to 50% hematocrit in Cytomix. 50 μg of linearized pMG75 MPPα plasmid isolated using a Qiagen Maxikit (Qiagen, Valencia, CA, USA) and precipitated in ethanol was centrifuged down and resuspended in 100 μl Cytomix. The plasmid and 250 μl of the parasite cytomix suspension were combined and transferred to a 0.2 cm electroporation cuvette on ice. Electroporations were carried out using a BioRad GenePulser set at 0.31kV, 960uF. The electroporated cells were immediately transferred to a T-25 flask containing 0.2 ml uninfected 50% RBCs and 10 ml medium. Parasites were selected using 2.5 µg/ml Blasticidin and 250 nM anhydrotetracycline (aTc).

### Transfection vector generation

Optimal gRNA sequences were identified using CRISPOR software to generate a double strand break within the endogenous MPP*α* locus (25). gRNA sequences were cloned into a Cas9-sgRNA vector (26)(S1A and B).

pMG75 (27) plasmid backbone was used to construct the repair template vector used to help generate D10_VP2KO_MPP*α*, the MPP*α* conditional knockdown line in the D10_VP2KO background. (24). A synthetic gene block was ordered from GenScript containing 584 base pair MPP*α* 5’ homologous region (nucleotide 1591-2175) in the coding sequence, containing silent blocking mutations in the selected gRNA recognition site, and 550 base pair 3’homologous region (nucleotide 2178-2728) in the 3’UTR, separated by a linker region containing restriction enzyme sites for SacII and BspEII for linearization of the plasmid. The homologous regions are flanked by BstEII and BssHII for ligation into the pMG75 backbone, which contains the TetR-DOZI-aptamer regulatory system, as well as a BSD selectable marker. The synthetic gene was inserted into the pMG75 plasmid backbone via T4 DNA ligase. All plasmids were sequenced using Plasmidsaurus (South San Francisco) nanopore sequencing. The plasmid was linearized with SacII and BspEII before transfection.

### Growth Assay

Parasites were enriched for late trophozoite and early schizont stages using a layered step gradient of 70%, 60%, and 35% Percoll. Equal amounts of synchronized parasites were split into a 6 well plate containing Human (O+) RBC at 2.5% hematocrit in 10ml of complete RPMI medium. Triplicate cultures were grown in the presence or absence of aTc, with daily medium change. Cumulative parasitemias were counted over the span of 8 days using flow cytometry after staining with SYBR Green. Cumulative parasitemias were plotted using GraphPad Prism.

### Immunofluorescence assays

50 μL of packed volume parasite cultures were fixed with 4% v/v paraformaldehyde and 0.0075% v/v glutaraldehyde with slow mixing in a rotor for one hour at room temperature. The cells were washed 3x with PBS, followed by permeabilization with 0.125% v/v Triton X-100 in PBS. The cells were treated with NaBH_4_ (0.1 mg/ml) for five minutes at room temperature while mixing on a rotor. Samples were washed twice with 1X PBS and then blocked with 3% w/v BSA for 1 hour. Anti-HA (Lot#12923, Santa Cruz Biotechnology) and anti HSP60 (Lot# PP487392 Novus Bio) antibodies were added to the parasites at 1:500 dilution, and the samples were incubated overnight at 4°C in a rotor. Following three PBS washes, Alexa Fluor 488 conjugated goat anti-mouse IgG (Invitrogen Lot# SG251135) and Alexa Flour 568 conjugated goat anti-rabbit (Invitrogen lot #2447870) secondary antibodies were added at 1:250 dilution, followed by overnight incubation at 4°C. After three PBS washes, the treated parasite samples were resuspended in anti-fade buffer (S2828, ThermoFisher Scientific) and imaged using a Nikon Ti microscope.

### Live Cell Microscopy

35 mm glass bottom culture dishes were coated with 0.1% poly-D-lysine overnight and washed 3 times with PBS. 250⍰μL of 2.5% hematocrit parasite culture was added to the dishes and incubated for 30 min to allow attachment. Culture dishes were washed 3 times with 1XPBS followed by addition of phenol red-free RPMI. Parasites were stained with 62.5 nM MitoTracker Red (Life Technologies by ThermoFisher Scientific) for 20 minutes, followed by 0.5 mg/ml Hoechst nuclear staining for 10min. Imaging was done using the Nikon Ti microscope. Parasites were temperature controlled at 37 ⍰C. Hoechst-stained parasites were visualized with the DAPI filter set, while MitoTracker red was visualized using the tetramethyl rhodamine isocyanate (TRITC) filter set. Images were 2-dimentionally deconvoluted and processed using Nikon NIS Elements (5.30.02) Imaging Software. The presence of mitochondrial membrane potential or parasite plasma membrane potential was visualized following incubation with MitoTracker red. For statistical analysis of the membrane potential visualization, 30 parasites were counted and imaged. The results were graphed using GraphPad Prism.

### Western blotting

Saponin lysed parasite pellets were suspended in approximately ten volumes of 2% w/v SDS, 5% β-mercaptoethanol, and 1% bromophenol blue. Samples were centrifuged for 5 minutes at 13,000g, and the supernatant was subjected to SDS-PAGE. The separated proteins on the gel were transferred to PVDM membranes at 20 volts overnight. Powdered milk solution was used to block the membranes. Blots were then incubated at a dilution ratio of 1:10,000 with primary antibody. After probing with primary antibody (Mouse monoclonal anti-HA Lot#12923, Santa Cruz Biotechnology, Rabbit anti- *Plasmodium* aldolase Lot#GR289448-4 ABCAM, Rabbit anti HSP60 Lot# PP487392 Novus Bio) blots were washed with Tris-buffered saline with tween (TBST) and incubated with secondary antibody (Goat anti mouse IgG-HRP Lot#GR3219575-4 ABCAM, Mouse anti rabbit IgG-HRP Lot#L1819 Santa Cruz Biotechnology) conjugated with horseradish peroxidase at a concentration of 1:10,000.

### Immunopulldown and Proteomics

100 μL parasite pellets from both wildtype and D10_VP2KO_MPP*α* were solubilized overnight at 4° C in solubilization buffer containing 2% digitonin, 200 mM 6-aminocaproic acid, 50 mM Bis-tris, pH 7, 1 mM EDTA, 1 mM AEBSF, and protease inhibitor cocktail (Sigma-Aldrich). The solubilized sample was centrifuged at 14,000 rpm for 15 min at 4 °C to remove debris, and the supernatant was incubated with anti HA antibody coated magnetic Dynabeads (Pierce Anti-HA, Clone 2-2.2.14, Clone 9E10; Thermo Fisher Scientific). Beads were washed 3 times in 1x PBS, and solubilized total protein was added to 50 μl of the beads, followed by overnight mixing by rotation at 4 °C. Protein-bound beads were washed three times with 1x PBS + 0.02% digitonin. Beads with bound proteins were used for LC/MS on bead proteomics at the University of South Florida Mass Spectrometry & Proteomic Core Facility. Protein samples were prepared for LC-MS/MS using S-Traps (Protifi). 5% SDS in 50 mM triethylammonium bicarbonate (TEAB) was added to the IP beads. Protein samples were reduced with 20 mM dithiothreitol (DTT) at 95 °C for 10 min and alkylated using 40 mM iodoacetamide (IAA) in the dark at room temperature for 30 min. The samples were acidified with 12% phosphoric acid (final concentration 1.2%) before adding 6x volumes of S-trap buffer (90% Methanol, 100 mM TEAB). The protein solution was loaded onto micro S-traps, and centrifuged at 4000xg for 30 sec. Three washes with 150 µl S-trap buffer were performed. 0.5 µg of Trypsin/LysC (Promega) was added to the S-trap filter and digested overnight (∼16 hrs) at 37 °C. Peptides were eluted with sequential additions of 50 mM TEAB, water+0.1% formic acid, and 50/50 acetonitrile/water+0.1% formic acid, centrifuged at 4000xg for 1 min each. Peptides were completely dried in a vacuum-centrifuge concentrator before resuspension in water+0.1% formic acid.

Peptides were separated using a Vanquish Neo UHPLC (Thermo) with a 50 cm EASY-Spray PepMap Neo C18 column (Thermo) and analyzed on a Q Exactive Plus hybrid quadrupole mass spectrometer (Thermo). Peptides were separated over a 1hr gradient (Mobile Phase A: water+0.1% Formic Acid, Mobile Phase B: Acetonitrile+0.1% Formic Acid; 2% B-32% B), and data was acquired using data-dependent acquisition (DDA) with a Top 30 method. Full MS scans were acquired at 70k resolution, MS/MS scans were acquired at 17.5k resolution. Raw data files were searched using MaxQuant (v2.4.9) against the current *Plasmodium falciparum* proteome sequences downloaded from PlasmoDB.org, as well as MaxQuant generated databases of reverse protein sequences and common contaminant protein sequences. Digestion settings specified Trypsin/P specific enzyme digestion, with a maximum of 2 missed cleavages and a minimum peptide length of 7. Both PSM (peptide spectral match) and protein FDR cutoffs were set at 1%. Cysteine carbamidomethylation was set as a fixed modification. Methionine oxidation and acetylation (protein N-term) were set as variable modifications. The MS/MS match mass tolerance was set at 20ppm. Label Free Quantification intensities of enriched proteins were compared across the replicates to determine fold change and statistical significance using pair wise Student t-test. Volcano plots were visualized using Graphpad Prism.

### Growth inhibition assay

All parasite growth inhibition assays were performed in triplicate in 96-well plates as described by Desjardin et al. (28). *P. falciparum*-infected erythrocytes had aTc removed 48 hours prior to the start of the assay. At 1.0% initial parasitemia and 1.5% hematocrit, these parasites were exposed to various concentrations of the indicated drug/inhibitor for 48h and then pulsed with 0.5 μCi of ^3^H-hypoxanthine for an additional 24h. Untreated parasitized and unparasitized red blood cells were incubated concurrently with the treated parasites as controls for growth and radioactive precursor incorporation. The 96 well plates were then frozen at −80°C for 24 h to disrupt erythrocyte and parasite membrane integrity. Following freezing, the plates were warmed to 37°C, and parasites were harvested onto EasyTab^™^ -C Self-Aligning Glass Fiber Filters (Packard, Meridian, CT, USA). The filters were then dried completely and placed into an Omnifilter™ 96-well plate filter case (Packard, Meridian, CT, USA). 30μl of OmniScint (Packard, Meridian, CT, USA) scintillation fluid was added to each well and the radiation was counted with a TopCount^™^ counter. The incorporation of radioactivity into nucleic acids served as a measure of cell proliferation and is reported by TopCount as counts per minute (cpm). The total cpm of the non-parasitized red blood cells was subtracted from the cpm of all other wells. The cpm of each sample was converted to percent by dividing it by the cpm of the untreated control. The percent growth versus drug/inhibitor concentration was plotted using GraphPad Prism software. A nonlinear regression dose-response curve was calculated in Prism after adjusting the lowest percent to equal zero and the highest percent to 100.

## Results

### Generation of parasite lines conditionally expressing MPPα

A parasite line expressing yDHOD (D10_VP2KO) was chosen as the parental line to prepare the MPP*α* conditional knockdown line. These parasites are independent of mtETC activity and are resistant to all mtETC inhibitors (5, 29). We modified the endogenous MPP*α* locus (PF3D7_0523100) in D10_VP2KO parasites using CRISPR-Cas9 to generate parasites conditionally expressing 3HA-tagged MPP*α* under regulation of the aptamer-TetR-DOZI system (Figure S1). PCR confirmed the integration of the expected portion of the donor plasmid within the modified MPP*α* locus (Figure 1A and B). Localization of MPP*α* -3HA was determined by immunofluorescence using anti-HA antibody (Figure 1C). We used HSP-60 antibody as a mitochondrial marker to demonstrate colocalization of MPP*α* at the parasite mitochondrion, as shown in Fig. 1C.

**Figure 1.**
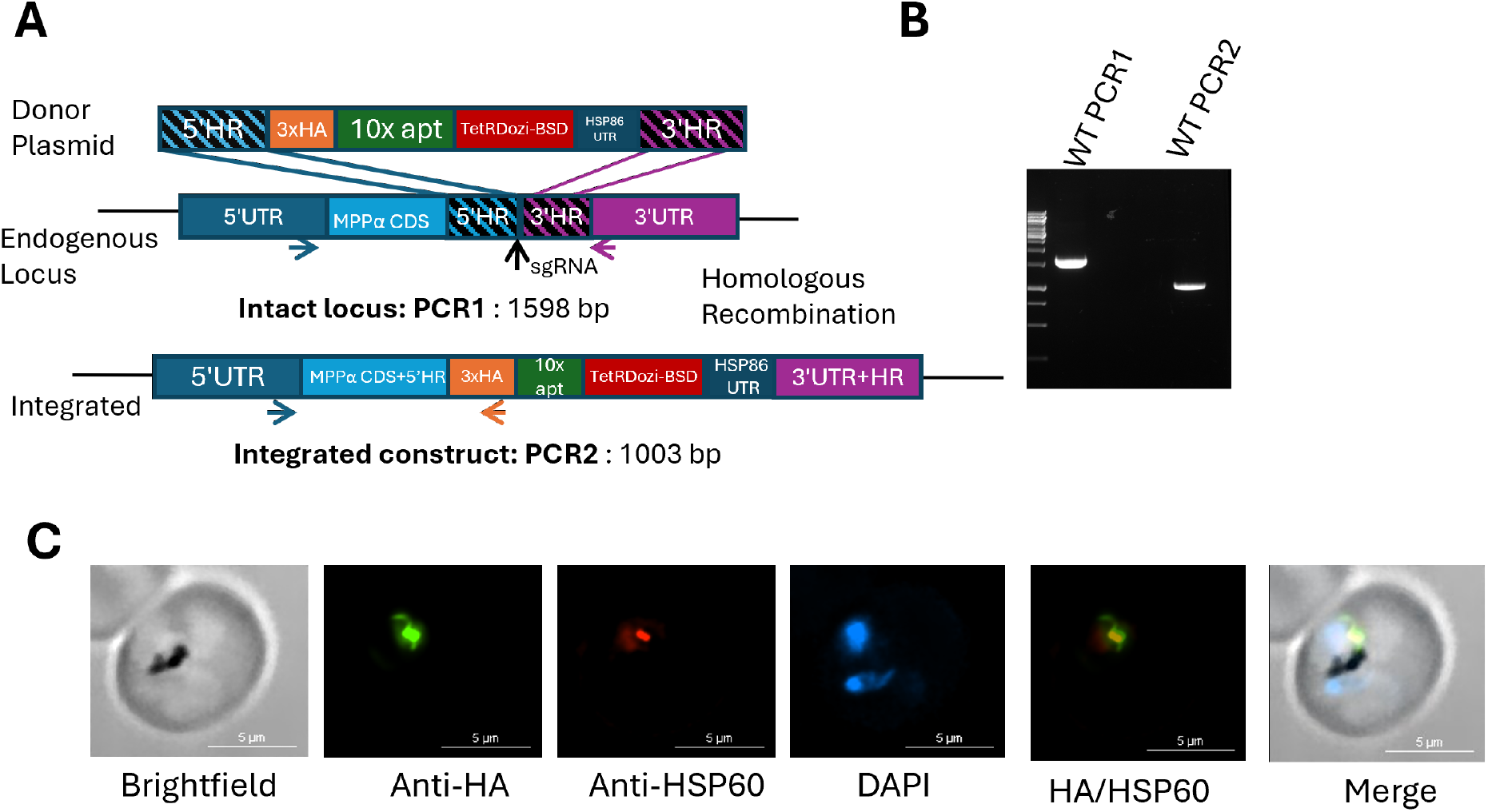
Generation of parasite lines conditionally expressing MPPα. (A) Schematic of MPP*α* locus modification using CRISPR-Cas9. Vertical arrow indicates position of sgRNA binding. Also indicated are the 5’ and 3’ homologous regions (HR), 5’ HR coming from MPP*α* CDS and the 3’ HR coming from 3’UTR. (B) Using primers indicated by PCR1, amplification of the forward and reverse primers from the endogenous locus shows that the locus is disrupted in the MPP*α* transfected line. Using primers indicated for PCR2, amplification of the forward primer from the endogenous locus and the reverse primer from the HA tag present in the plasmid determines correct integration of the donor DNA into the parasite genome. (C) Representative immunofluorescence images of a parasite conditionally expressing MPP*α*, showing its mitochondrial localization.

### MPPα is essential for parasite viability

We assessed expression of MPPα in the presence and absence of aTc by carrying out western blot analysis of parasite lysates collected over 8 days of parasite cultures; HSP60 expression was used as a loading control. As shown in Fig. 2A and B, expression of MPP*α* was below detection level by day 4 of parasite growth in absence of aTc. We assessed the effects of MPP*α* knockdown on parasite growth by removing aTc at the trophozoite stage of the parasite. As shown in Fig. 2C, parasite growth ceased in the second cycle following aTc removal. No viable parasites were detected via Giemsa stain in MPP*α* (-) condition by day 6 following aTc removal. Since these studies were carried out using a parasite line that was independent of mtETC activity by having the metabolic bypass provided by yDHOD, suggesting that MPPα serves a critical function in addition to supporting mtETC activity as a subunit of complex III.

**Figure 2.**
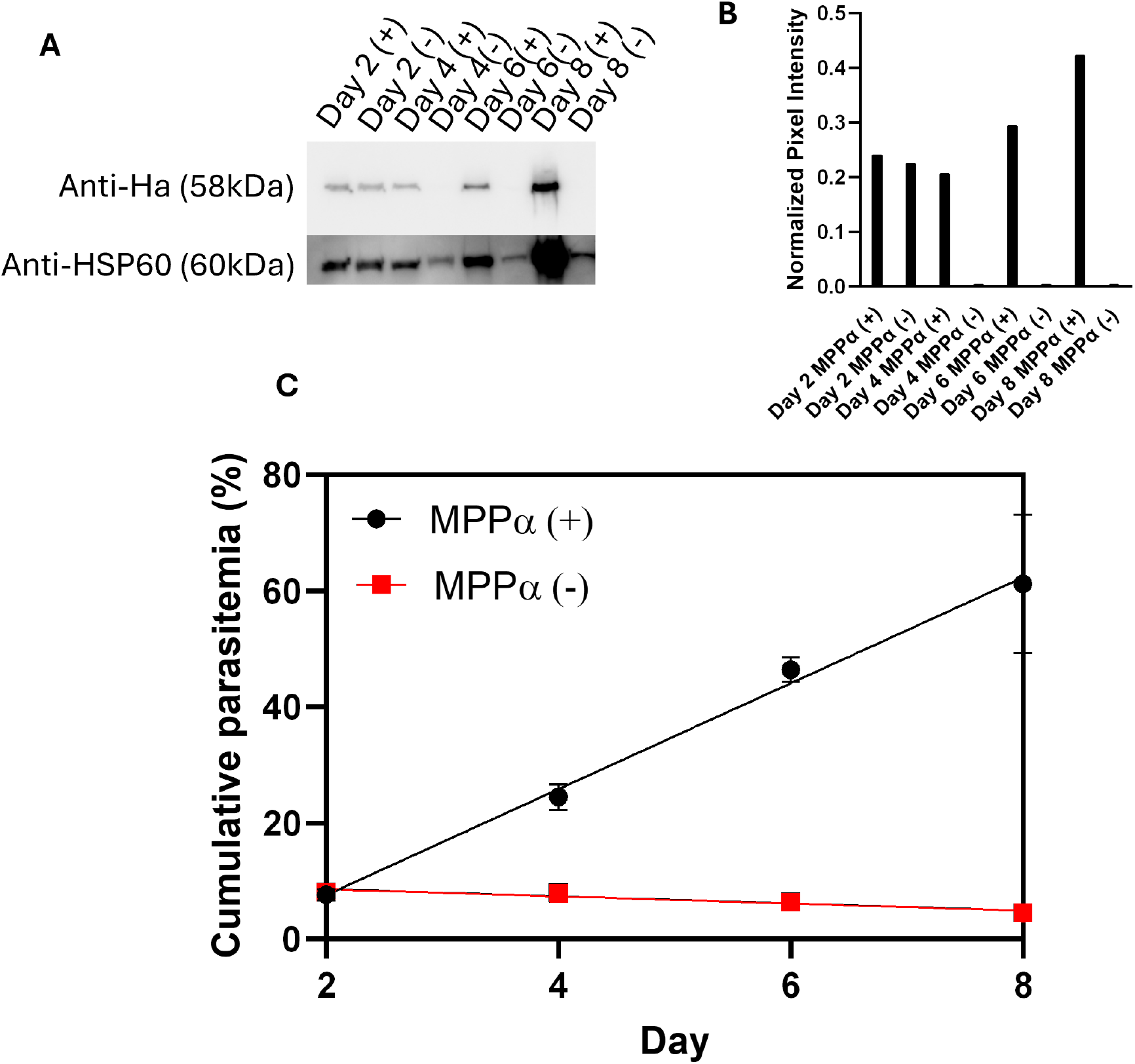
MPPα is essential for parasite viability. (A) Western blot analysis using anti-HA and anti-HSP60 antibodies. (B) Quantification of Western blot normalized to intensities of anti-HSP60. Pixel intensities were measured via Image J.⍰(C) Assessment of growth of MPPα (+) and MPPα(-) parasites. Growth assay data was conducted in triplicate by flow cytometry using SYBR green stained parasites.

### Mitochondrial membrane potential is disrupted in knockdown parasites

Maintenance of electrochemical potential across the inner mitochondrial membrane is critical for the survival of all eukaryotes. We have previously shown that blood stages of *Plasmodium falciparum* have two independent means to maintain mitochondrial membrane potential: one dependent on mtETC and the other independent of mtETC (5). Therefore, we next examined mitochondrial membrane potential in parasites with and without MPPα expression. Using live cell microscopy, the presence of mitochondrial membrane potential was visualized using Mitotracker red staining in both knockdown and wild type parasites. After one cycle of MPPα knockdown, 40% of the parasites were observed to have a diffused Mitotracker staining, which is indicative of collapsed mitochondrial membrane potential. Since parasite plasma membrane potential remains intact (initially), the positively charged probe accumulates within the cytoplasm, but dissipation of the mitochondrial membrane potential would prevent its accumulation in the mitochondrion. After two cycles, 73% of knockdown parasites displayed diffuse staining. By Day 6 of MPPα knockdown, no parasites with mitochondrial staining were visible (Figure 3A and B). While MPPα depletion may directly affect mtETC, the secondary pathway for mitochondrial membrane potential should remain operative. Thus, these data indicate that removal of MPPα affects both the primary and secondary pathways for generating membrane potential.

**Figure 3.**
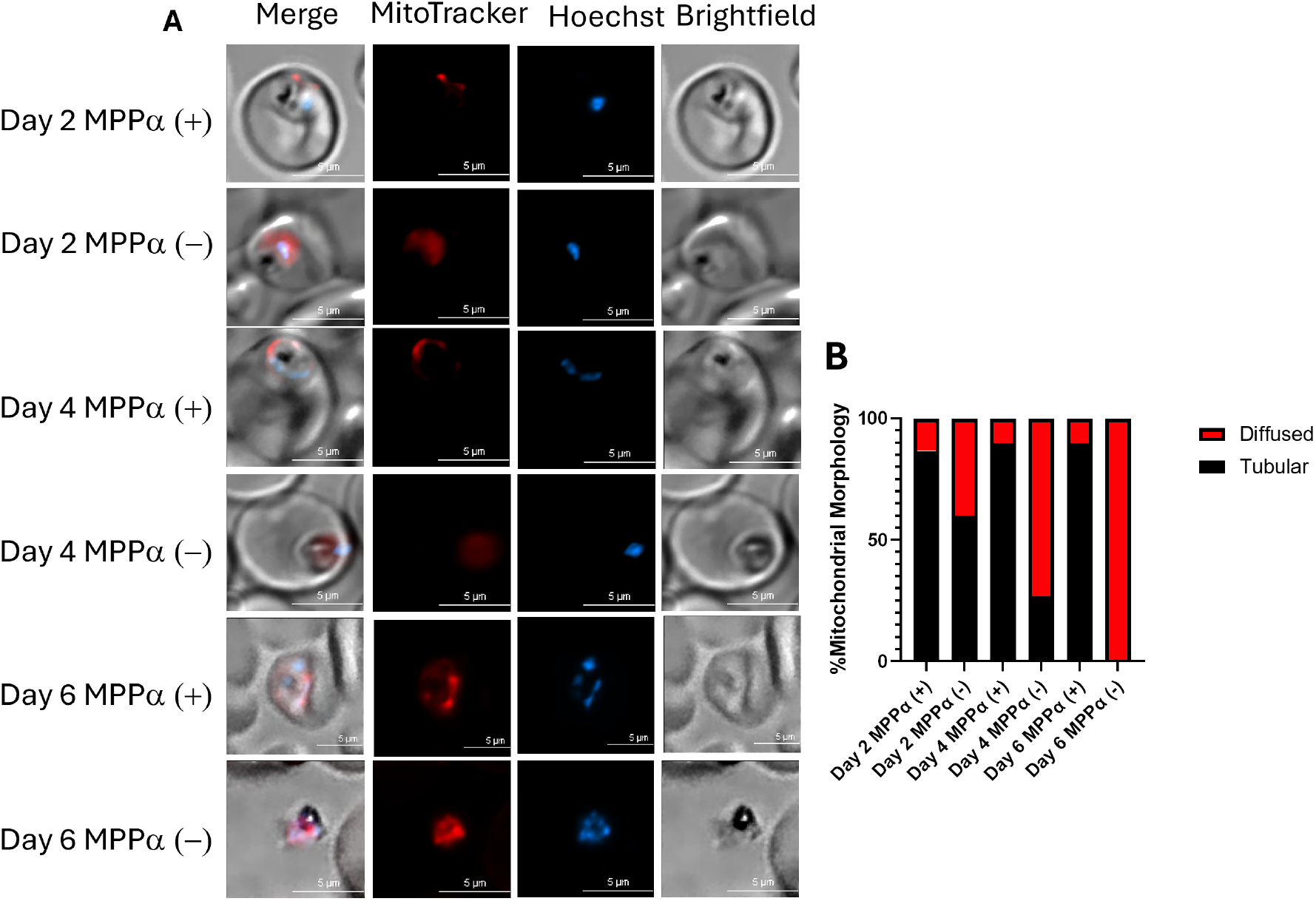
Mitochondrial membrane potential is disrupted in knockdown parasites. (A) Live cell microscopy images of MPPα (+) and MPPα (–) parasites. Mitotracker staining (red) revealed tubular morphology in MPPα(+) parasites but was diffused in varying number of MPPα(-) parasites starting as early as Day 2 following the knockdown (B) Quantitation of morphological profiles of 30 parasites at each time points was assessed, showing gradual increase in parasites showing diffuse Mitotracker staining resulting from disruption of mitochondrial membrane potential.

### Knockdown of MPPα expression causes hypersensitivity to proguanil

A commonly used antimalarial drug, Malarone, is a synergistic combination of atovaquone and proguanil (30, 31). We have previously shown that the synergistic property of proguanil results from its inhibition of the secondary pathway for mitochondrial membrane potential generation (5). Since MPPα depletion affects both mtETC dependent and independent pathways of generating membrane potential (Figure 4A), we wanted to investigate whether the absence of MPPα influences proguanil sensitivity of the parasites. We carried out growth inhibition assays using ^3^H-hypoxanthine incorporation by parasites treated with proguanil in presence and absence of MPPα. While MPPα(+) parasites showed EC50 of 1585 nM for proguanil, MPPα(-) parasites were approximately 200-fold hypersensitive to proguanil with EC50 of 8.4 nM (Figure 4B). In contrast, there were no changes in response of the parasites to other inhibitors (DSM1, atovaquone and PA21A092) in presence or absence of MPPα (Figure S3). We have previously shown that ubiquinone analogue, decyl-ubiquinone, is able to partially rescue parasite growth when subjected to Complex III inhibition (32). Thus, we assessed the ability of decyl-ubiquinone to reverse hypersensitivity to proguanil in MPPα(-) parasites. Addition of 25 *μ*M decyl-ubiquinone had only a relatively minor effect on hypersensitivity to proguanil in MPPα(-) parasites (Figure 4C).

**Figure 4.**
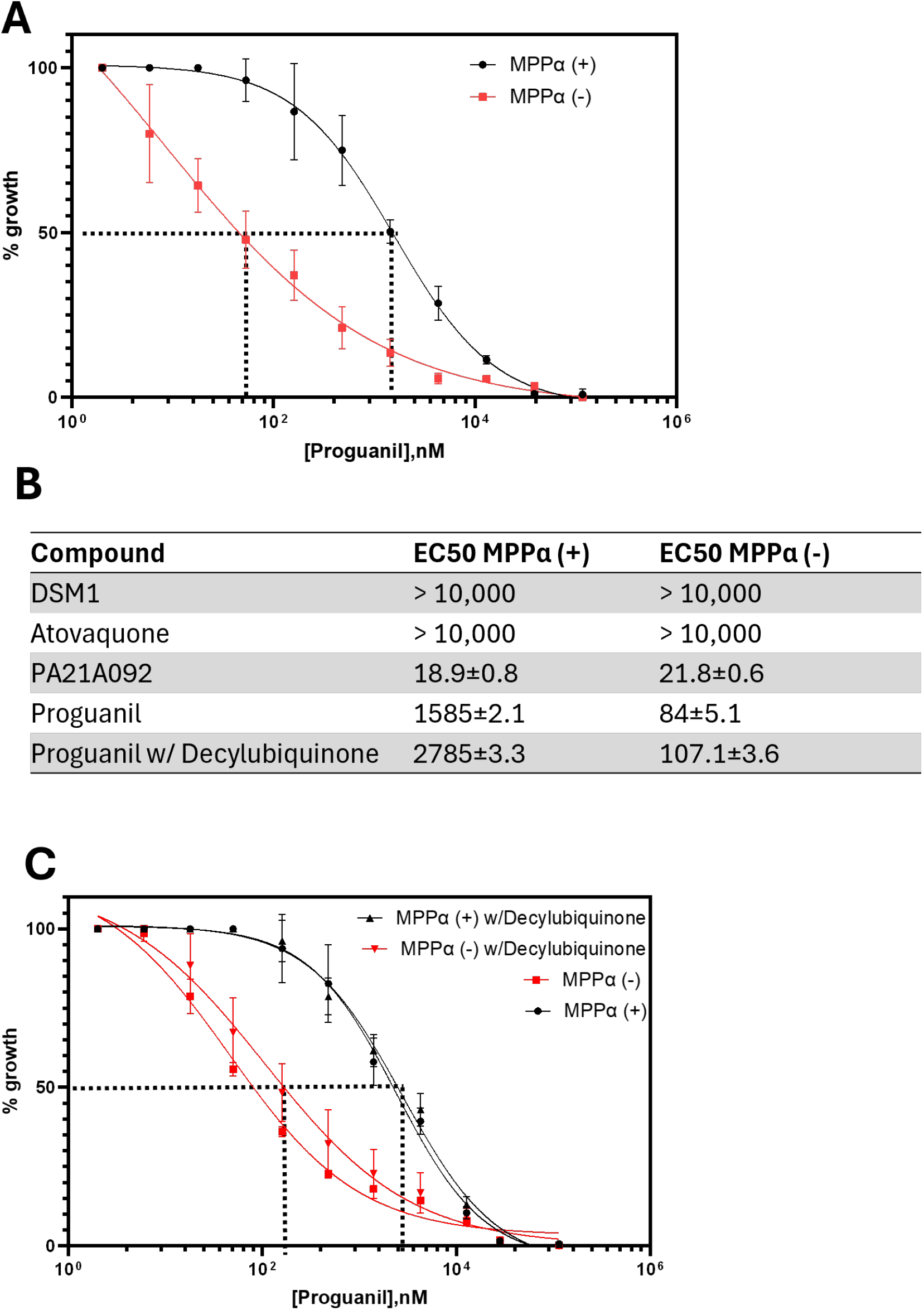
Knockdown of MPP*α* expression causes hypersensitivity to proguanil. (A) Parasite growth inhibition assessed by ^3^H-hypoxanthine incorporation following proguanil treatment. For MPP*α*- parasites, aTc was removed for 48 h followed by ^3^H-hypoxanthine addition for 72hr. MPP*α*(+) parasites were also grown for 72hr followed by ^3^H-hypoxanthine addition. (B) EC_50_ values for DSM1, atovaquone, PA21A092, and proguanil in MPP*α*+ and MPP*α*- parasites. While the EC50 values for Complex III, DHOD and PfATP4 inhibitors were similar for MPP*α*(+) and MPP*α*(-) parasites, MPP*α*(-) parasites were almost 200-fold hypersensitive to proguanil. (C) Parasite growth inhibition assessed by ^3^H-hypoxanthine incorporation following proguanil treatment and addition of 25 *μ*M decyl-ubiquinone where indicated. For MPP*α*(-) parasites, aTc was removed for 48 h followed by H-hypoxanthine addition for 72hr.

### Proteomic analysis of MPP*α* immunoprecipitation

To gain insight into interactions of MPP*α* with other mitochondrial proteins, we carried out immunoprecipitation from both MPP*α* tagged and wildtype parasites using anti-HA beads followed by LC-MS/MS analysis. Overall, 93 proteins were identified as significantly enriched in the MPP*α* immunoprecipitated protein samples, using a cutoff of two-fold change compared to the control with p-value <0.05 (Figure 5A). Of the putative 12 subunits of Complex III, 11 subunits, including mitochondrially encoded cytochrome *b*, were enriched greater than a million-fold, based on the label free quantification (LFQ) intensity of the peptides observed. QCR13 (Pf3D7_0817800) was the only putative Complex III protein that was absent, likely due to its small size of just 54 amino acids. Remarkably, there were 45 additional predicted or experimentally confirmed mitochondrial proteins that were equally significantly enriched. This suggests their interactions with MPP*α* are of similar magnitude as MPP*α*’s association with Complex III. We postulate the presence of these mitochondrial proteins with MPP*α* being by virtue of their transient association during processing of their mitochondrial targeting peptides. The significance of additional proteins that were enriched but not predicted to be mitochondrial remains unclear. It is interesting to note, that cytochrome *c* oxidases subunits 1 and 3 were not enriched. This is likely due to these proteins being encoded by mtDNA and not requiring processing of mitochondrial targeting sequences (1,6) . These data suggest that Complex III is engaged in the dual functions of mtETC integrity, as well as processing imported mitochondrial proteins.

**Figure 5.**
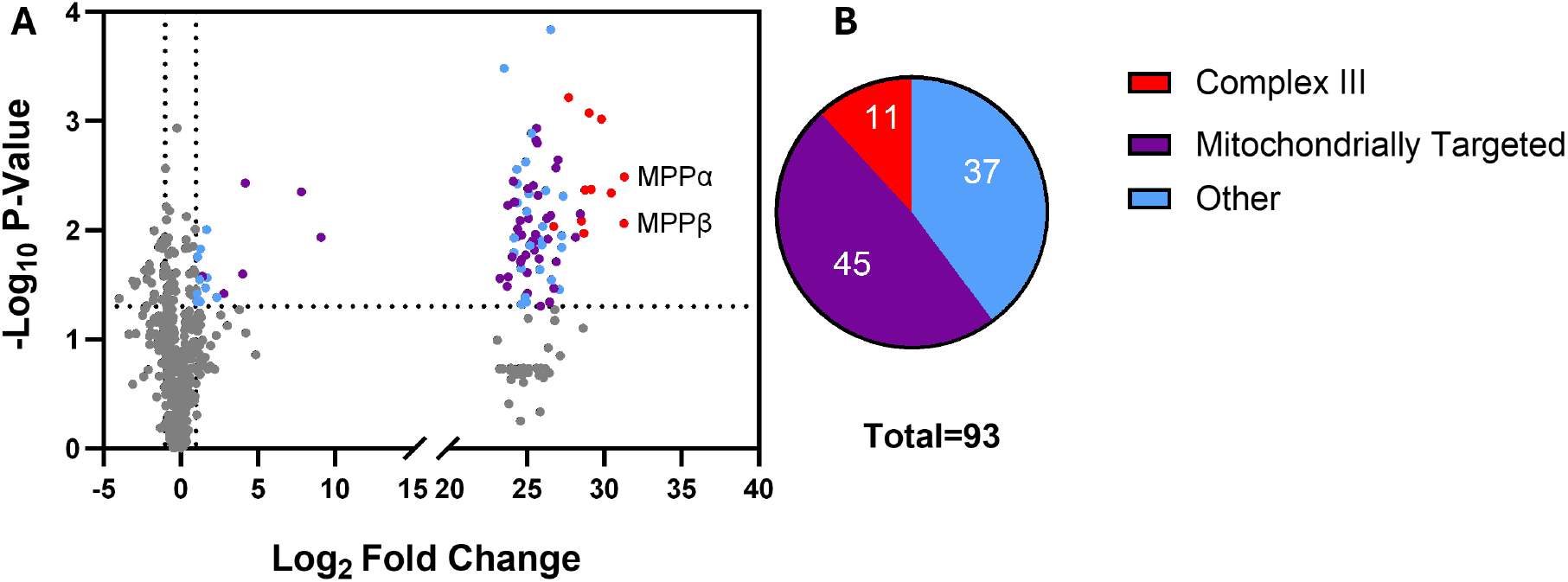
Proteomic analysis of MPP*α* immunoprecipitation. (A) Volcano plot of enriched proteins with a log_2_-fold change of label free quantification (LFQ) intensity and p-values of < 0.05 found in immunoprecipitated MPP*α* samples. The red dots represent proteins present in Complex III. The purple dots represent proteins that are predicted to be targeted to the mitochondria. The blue dots represent the other proteins that were significantly enriched in the immunoprecipitated samples. (B) Pie chart depicting the number of proteins that fall into the categories of Complex III, likely mitochondrially targeted, and other enriched proteins.

## Discussion

Although mitochondria in all extant organisms originated from the symbiotic arrangement between an archaeon and an *α*-proteobacterium that led to the emergence of eukaryotes, mitochondrial genomes and physiology are as divergent as the organisms in which they reside (33, 34). Oxidative phosphorylation to generate ATP, while a major function of mitochondria in many organisms, is not essential in some organisms, especially those that live in anaerobic environments (35). Indeed, such organisms often lack mtDNA but continue to possess a remnant mitochondrion called a mitosome (36, 37). In addition, certain yeasts (38), cultured vertebrate cells (39) and bloodstream-forms of trypanosomes (40) can be manipulated to lose their mtDNA while maintaining the mitochondrion itself. These are called rho^0^ cells, and investigations with such rho^0^ cells have helped to establish that the absolute essential function of their retained mtDNA-negative mitochondria is to synthesize and export inorganic Fe-S clusters for a variety of iron-sulfur proteins required for cell survival (41, 42). To permit this essential function, mitochondria need to maintain an electropotential across their inner membrane, which is required for protein and metabolite transport. In mitochondria with an active mtETC the membrane potential is normally established through proton pumping by the respiratory complexes. A lack of mtDNA will lead to the absence of mtETC, so then the electropotential will need to be generated through alternative means, generally by reverse action of ATP synthase and/or by electrogenic exchange of ATP/ADP (43).

Asexual blood stage *P. falciparum* rely on glycolysis for ATP generation with little contribution from mitochondrial oxidative phosphorylation (44, 45). Yet the parasite maintains an active mtETC, inhibition of which by antimalarials such as atovaquone results in parasite death. We have previously shown that the critical function of mtETC in blood stage *P. falciparum* is to re-oxidize ubiquinol to support mitochondrial DHOD, which is needed for pyrimidine biosynthesis, an essential pathway in the parasites (5). Parasites engineered to bypass mitochondrial DHOD become mtETC independent and resistant to drugs such as atovaquone. We also showed that generation of electropotential across the mitochondrial inner membrane was still required in these mtETC-independent parasites, and that the generation of this potential could be inhibited by proguanil, resulting in parasite death when combined with mtETC inhibitors (5). Since mtDNA in *P. falciparum* encodes just three components of mtETC, we initially reasoned that it should be possible generate rho^0^ parasites. However, various attempts failed to generate such parasites (8-10). It is of interest to note that all organisms where rho^0^ cells can be generated contain matrix-localized MPP (e.g., yeast, trypanosomes, and mammals), whereas organisms with Complex III associated MPP, such as plants (and *P. falciparum*), cannot be rendered rho^0^. Because cytochrome *b* of Complex III is always encoded by mtDNA, maintenance of mtDNA remains essential in these organisms for proper assembly of MPP within Complex III, even when mtETC is made non-essential.

A previous study by Espino-Sanchez et al. (46) assessed effects of conditionally knocking down expression of cytochrome *c*_*1*_, a component of Complex III, in *P. falciparum*. They showed that the parasites lacking this component of Complex III failed to survive but could be rescued by addition of decyl-ubiquinone, consistent with Complex III being the source of ubiquinone to support DHOD. Furthermore, parasites lacking cytochrome *c*_*1*_ continued to maintain mitochondrial membrane potential, which could be collapsed by treatment with proguanil (46). These results are consistent with prior inferences that Complex III is essential for mtETC activity. In contrast to cytochrome *c*_*1*_ knockdown, however, we show here that knockdown of MPP*α* not only results in parasite demise but also a collapsed mitochondrial membrane potential. Parasites with MPP*α* knockdown cannot be rescued by either the presence of yDHOD or addition of decyl-ubiquinone. Since MPP*α* is a component of Complex III, our study establishes a secondary role of MPP in addition to maintaining the structure and function of Complex III of mtETC. We posit that this secondary role is to process mitochondrially targeted proteins. In support of this, we observed peptides from multiple mitochondrially targeted proteins in proteomic analyses of MPP*α* pulldown samples (Figure 5). We suggest that these peptides are generated from proteins transiently associated with Complex III as they are being processed by the MPP subunits.

## Acknowledgements

We thank Dr. Dale Chaput at University of South Florida Advanced Research Core for Mass Spectrometry for assistance with proteomic analysis. This work was supported by Grant number R01 AI028398 from National Institutes of Health.

S1: **Maps of plasmids used for transfections**. Maps of the two gRNA plasmids (A) and (B), and the map for double cross over modification of MPP*α* locus (C) are shown.

S2: **MPP*α* disruption in a second parasite line not expressing yDHOD**.(A) Assessment of growth of MPP*α* + and MPP*α* - parasites in Dd2attb parasites. Growth assay was conducted in triplicate by counting 500 parasites via Giemsa stain. (B) Western blot analysis using anti-HA and anti-Aldolase antibodies.

S3: **Susceptibility of MPP*α* - parasites to antimalarials remains unchanged**. Parasite growth inhibition assessed by ^3^H-hypoxanthine incorporation following DSM1, atovaquone, PA21A092 treatment. For MPP*α* - parasites, aTc was removed for 48 h followed by 3H-hypoxanthine addition for 72 hr.

