## Supplementary figures and images for "Dual Function of Mitochondrial Complex III in *Plasmodium falciparum*"

### Fig. S1

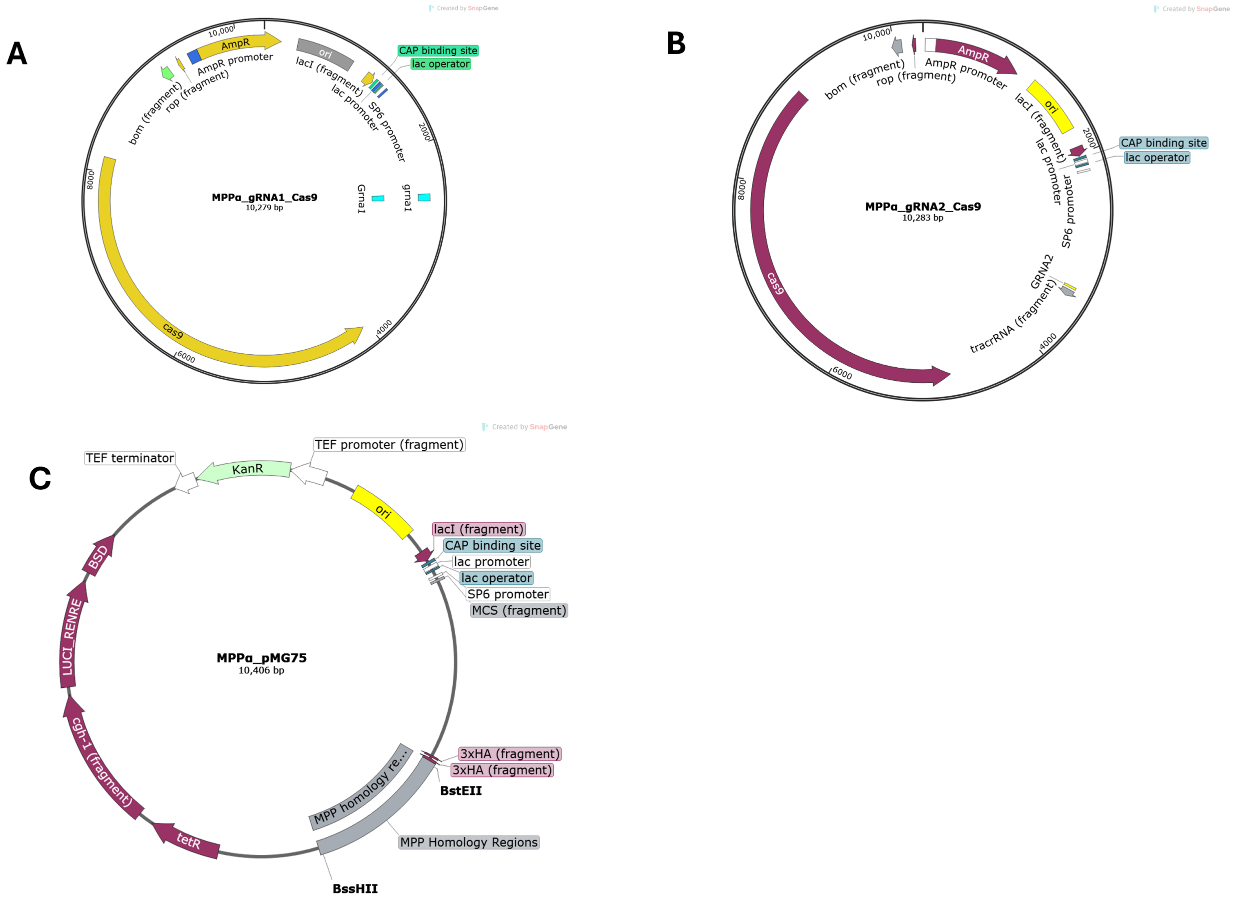

### Fig. S2

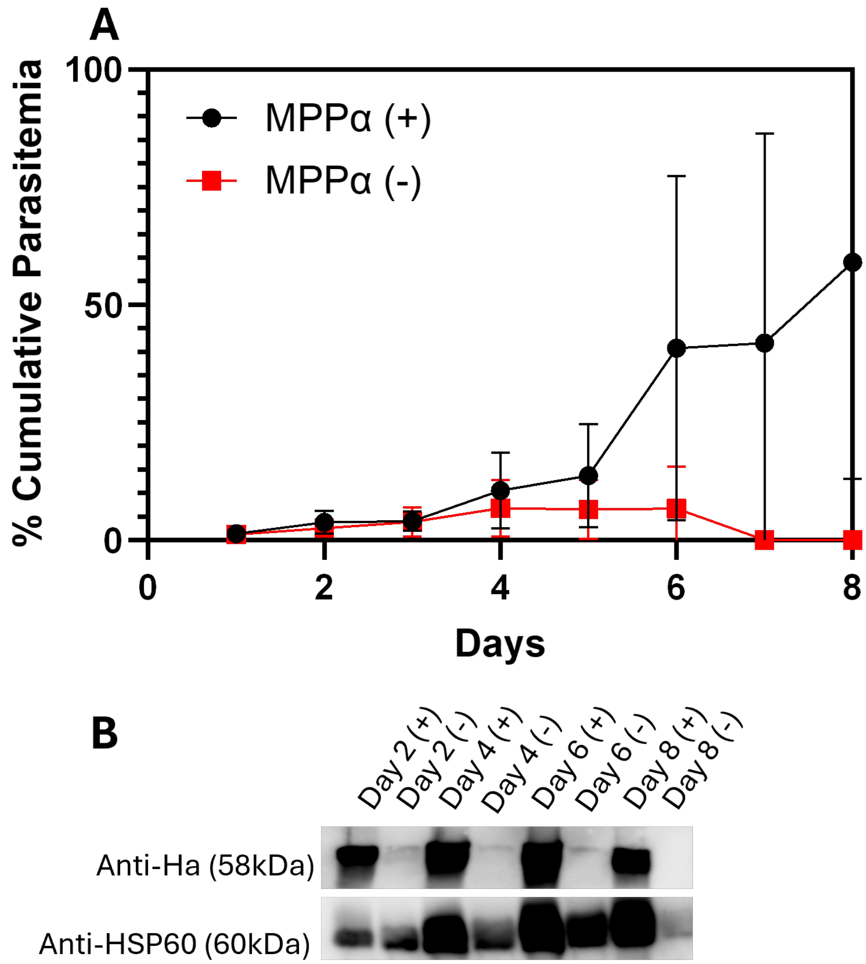

### Fig. S3

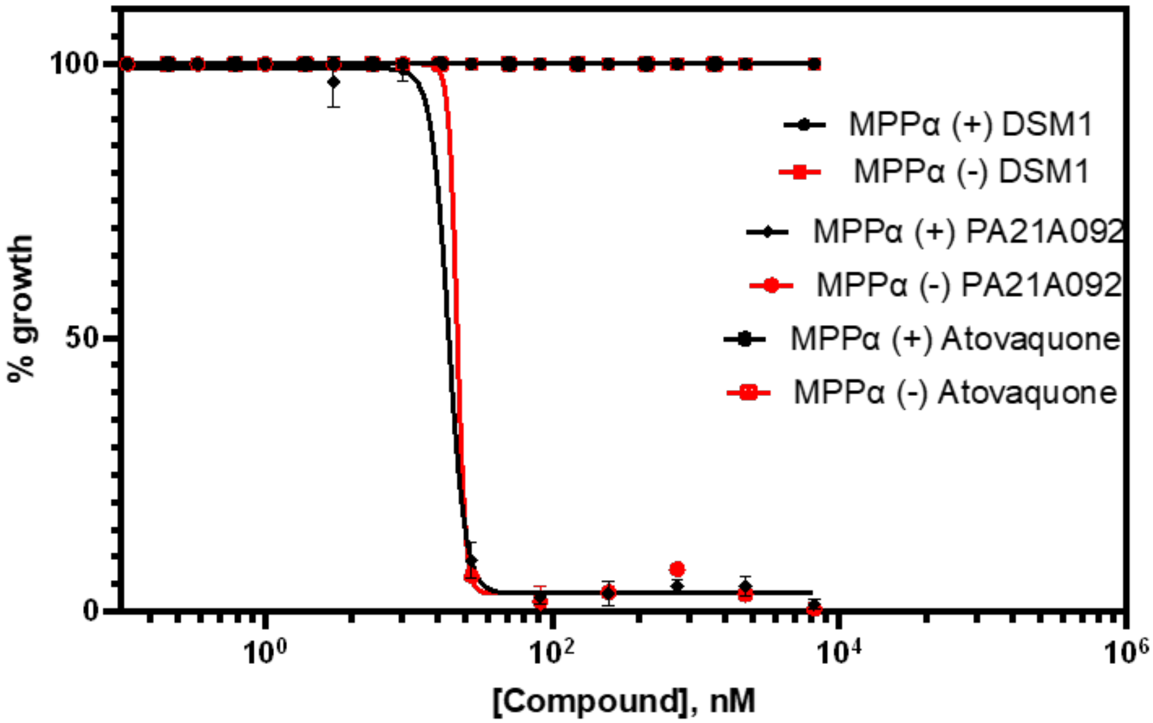
